# A closed-loop stimulation approach with real-time estimation of the instantaneous phase of neural oscillations by a Kalman filter

**DOI:** 10.1101/2021.04.25.441309

**Authors:** Takayuki Onojima, Keiichi Kitajo

**Affiliations:** CBS-TOYOTA Collaboration Center, RIKEN Center for Brain Science, Wako 351-0198, Japan; Division of Neural Dynamics, Department of System Neuroscience, National Institute for Physiological Sciences, National Institutes of Natural Sciences, Okazaki 444-8585, Japan; Department of Physiological Sciences, School of Life Science, The Graduate University for Advanced Studies (SOKENDAI), Okazaki 444-8585, Japan

**Keywords:** EEG, instantaneous phase estimation, real-time system, closed-loop, autoregressive model, Kalman filter

## Abstract

We propose a novel method to estimate the instantaneous oscillatory phase to implement a real-time system for closed-loop sensory stimulation in electroencephalography (EEG) experiments. The method uses Kalman filter-based prediction to estimate current and future EEG signals. We tested the performance of our method in a real-time situation. We demonstrate that the performance of our method shows higher accuracy in predicting the EEG phase than the conventional autoregressive model-based method. A Kalman filter allows us to easily estimate the instantaneous phase of EEG oscillations based on the automatically estimated autoregressive model implemented in a real-time signal processing machine. The proposed method has a potential for versatile applications targeting the modulation of EEG phase dynamics and the plasticity of brain networks in relation to perceptual or cognitive functions.

## 1. Introduction

Oscillations and synchronization are important features of neural activity that help us to understand how the brain implements perceptual and cognitive functions [1–3]. Synchronous neural oscillations are considered to contribute to various brain functions by mediating the communication between distant neuronal groups [4–7]. The development of analytic and experimental approaches, which can reveal the relationship between these neural dynamics and brain functions, is important in the field of neuroscience.

Neural oscillations can be recorded using electroencephalography (EEG), and the spontaneous EEG phase has been reported to influence the perceptual response to a stimulus. Previous studies have suggested that EEG phase dynamics mediate the crucial mechanisms to implement brain functions by modulating the sensitivity of neuronal responses or by dynamically changing the connectivity of global neuronal networks [8–11]. Methods to analyze the phase of EEG oscillations are, therefore, essential to reveal the functional role of rhythmic brain activity.

One of the more advanced approaches is closed-loop transcranial magnetic stimulation (TMS)-EEG [12], which has attracted much recent attention in neuroscience research [13–16]. This approach applies TMS over the brain to target the specific EEG phase. TMS is a non-invasive form of brain stimulation that induces an electric current at a specific area of the brain based on the principle of electromagnetic induction, with a temporal resolution of microseconds and a spatial resolution of millimeters [17, 18]. The closed-loop TMS-EEG approach has been reported to effectively induce plasticity in the motor cortex [16]; it has also been used to investigate the phase-dependent effect of evoked responses and could be used to reveal the role of neural oscillations for the implementation of cognitive and perceptual functions [13].

The present study focused on the phase-dependent stimulation approach [16, 19, 20], which has been applied to closed-loop TMS-EEG using autoregressive (AR) model-based prediction of EEG signals. This technique has the potential to extract information about the phase-dependent dynamical changes of global neural networks and to reveal the phase-dependent changes of plastic neural networks. The plasticity and dynamics of brain activity are induced by sensory and perceptual stimulation, such as visual or audio stimuli. However, the closed-loop approach has not been used in closed-loop EEG experiments using perceptual stimulation, due to various technical problems. This paper proposes a novel phase-dependent stimulation approach that was developed to avoid the following technical problems related to using a perceptual stimulus in the closed-loop EEG experiment.

A major technical problem of the closed-loop stimulation that prevents it from being applied to perceptual tasks is a decreased accuracy of the forward prediction of the EEG phase, which is caused by the longer time delay of the neural responses to perceptual stimuli compared with TMS-evoked neural responses. The phase-dependent stimulation approach using TMS needs to predict future EEG signals; one way to do this is to use AR model prediction to calculate the current and future EEG phase in real-time, and to directly apply stimulation to the brain at the targeted EEG phase. If sensory stimulation, instead of TMS, is used in a closed-loop system, it is necessary to predict longer-term EEG signals than in the case of direct brain stimulation by TMS because sensory stimulation has a longer time delay in reaching the cortex. The longer delay produces a decrease in the accuracy of phase-dependent stimulation. Therefore, it is necessary to develop an accurate real-time phase prediction method that has higher accuracy than the conventional method.

As another problem, the optimal parameters for the bandpass filtering and AR prediction need to be carefully chosen to accurately predict the EEG phase in the conventional AR model-based method. Notably, the conventional method has a trade-off between the order of the filter and prediction length, which affects the performance of EEG phase detection. This limitation makes it difficult to implement the closed-loop approach in EEG experiments. To do so, it is necessary to develop a novel method that is not almost necessary to select the value of parameters nervously by avoiding the trade-off problem.

In this study, we resolved these two technical problems by introducing the time-varying Kalman filter with an automatically estimated AR model for the phase-dependent stimulation method. To evaluate the accuracy of our method, we applied our method and the conventional method to eyes-closed resting EEG data with an offline analysis. As a result, we demonstrated that our proposed method can more accurately predict the EEG phase than the conventional method. Finally, we tested and implemented our method in real-time situations constructing the framework of the EEG replaying experiment. These results indicate that our method has advantages over the conventional method and is implementable in real-time situations.

## 2. Methods

### 2.1. EEG phase prediction by the AR model in a real-time situation

First, we will briefly explain the procedure of the conventional method, which uses the AR model to predict the EEG phase [16, 19–22]. In addition to predicting the EEG phase, the AR model has been applied to EEG signals to analyze the property of time-series data [23, 24]. The conventional method was used to make a comparison with our method, and to confirm the relationship between prediction performance and the alpha power of EEG oscillations (8–13z) in the eyes-closed resting condition.

The AR prediction algorithm was comprised of several sequential steps (Fig. 1a), as follows: 1) zero-phase bandpass filtering based on the target-frequency range, which was the alpha frequency range (8–13 Hz); 2) estimating the AR coefficients from the filtered signal; 3) predicting the future EEG signal using the AR model, and; 4) calculating the instantaneous phase of the analytic signal obtained using the Hilbert transform. The bandpass filter was used as an anti-causal two-pass finite impulse response bandpass filter (filter order 128) for the 8–13 Hz alpha band. After applying the bandpass filter, the edge data of the filtered signal (the range was 64 samples) were removed to avoid the edge effect caused by zero paddings. To predict the future signal, AR model coefficients were estimated. The AR model of order *m* is defined as follows:

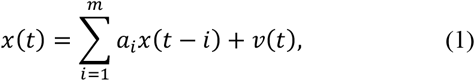

where *a*_1_,…, *a*_*m*_ were coefficients of the AR model, *x*(*t*) was a filtered EEG signal at time *t*, and *v* was white noise. The order was fixed as *m* = 30. The AR coefficients in this study were estimated using the Yule-Walker method. After estimating the AR coefficients, the prediction of future EEG signals was conducted based on the estimated AR model and filtered EEG signal. Finally, the EEG phase was calculated for the analytic signal of the predicted EEG signal obtained using the Hilbert transform [25].

**Figure 1.**
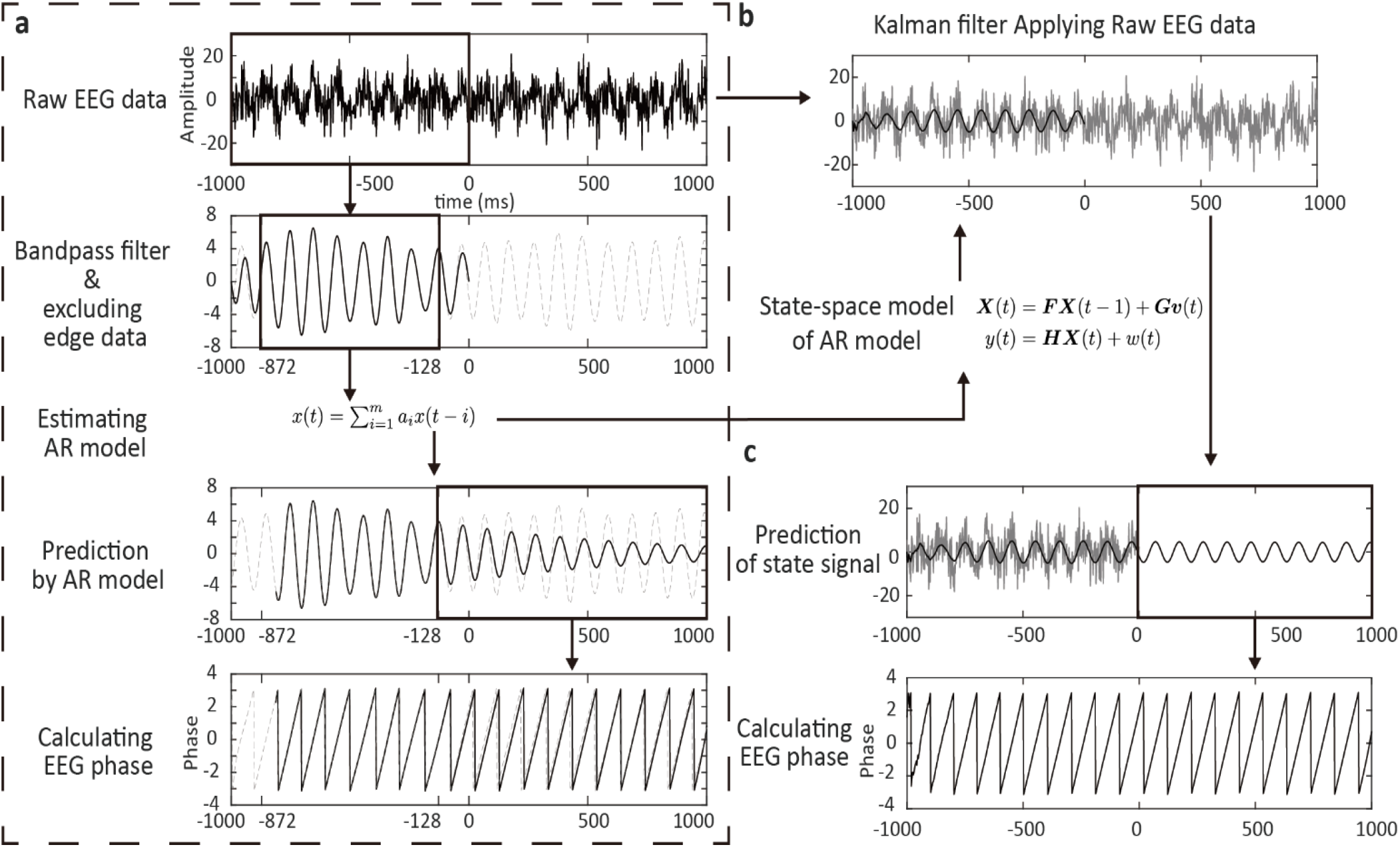
Schematic illustration of the conventional and Kalman filter-based methods. (a) The conventional method to detect the instantaneous phase. (b) Kalman filter-based AR model applied to raw EEG data. (c) Prediction based on the Kalman filter and calculation of the instantaneous phase.

The parameters to improve the prediction performance, such as the order of bandpass filter, the order of AR model, and the number of samples removed at the edge, were the same as those of a previous study [20]. We did not attempt to search for more optimal parameters because the aim was to investigate the benefits of introducing the Kalman filter. The parameters used in this study were chosen based on a previous study [16], and were as follows: the sampling rate was 500 Hz, the window size (the number of samples) was 500, the filter order was 128, and the number of samples removed at the edge was 64.

### 2.2. Estimating periodic neural activity using the Kalman filter

The conventional AR model prediction method is simple and practical for real-time implementation, and its performance largely depends on the results of the zero-phase bandpass filter [16]. If the filtered signal is contaminated by low-frequency signals or high-frequency noise, AR prediction and phase calculation become impossible. Therefore, it is necessary to accurately extract the target-frequency signal using a bandpass filter with a high filter order. However, there is a trade-off between the order of filters and prediction length because edge data must be removed to avoid the zero-padding effect and the length of removed edge data depends on the order of filters. If the order of the filter is sufficiently high to retain the filtering performance, it is necessary to predict a longer time. Therefore, the parameters of filtering and removing edge data should be chosen carefully. To resolve this issue, we improved the real-time prediction method by applying the Kalman filter.

We will now introduce the Kalman filter of an AR model. The following model is called the state-space model [26]:

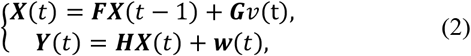

where ***X*** is the state vector; ***Y*** is the observation vector, which only has one dimension in this case; and *v* is the system noise, which is white noise with a mean of zero and variance *Q*. However, *w* is the observation noise, which is assumed to be white noise with a mean of zero and variance *R*. In our model, *y*(*t*) represents the raw EEG signals, *x*(*t*) represents the alpha EEG signal, and the state vector and observables are defined as ***X***(*t*) = (*x*(*t*), *x*(*t* − 1),…, *x*(*t* − *m* + 1))^*T*^, and ***Y***(*t*) = *y*(*t*), respectively. ***F***, ***G***, and ***H*** can be described by the following matrices using AR coefficients:

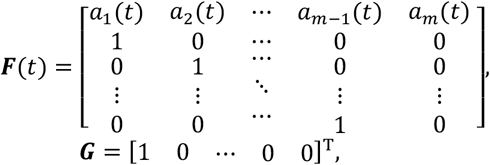

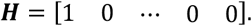

Here, ***F***(*t*), ***G***, and ***H*** are *m* × *m*, *m* × 1, and 1 × *m* matrices, respectively. The AR model coefficients *a*_*i*_ (*t*), which were components of the ***F***(*t*) matrix, and variance of system noise *Q*(*t*) were estimated using the Yule-Walker method based on the filtered signal (Fig. 1b). That is, ***F***(*t*) and *Q*(*t*) are decided based on the bandpass-filtered EEG signal for the past one second, except for the edge data, whose range represents the size of the window of the AR prediction method. Additionally, assuming that the filtered signal is generated from the state model, the variance of observed noise *R*(*t*) is calculated by obtaining the variance of the signal and subtracting the filtered signal from the raw EEG signal for the past second. The Kalman filtering and prediction of the AR model are easily implemented in real-time using the following equations:
[One-step-ahead prediction]

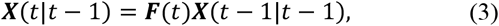

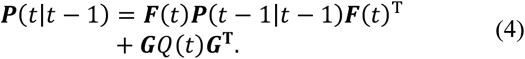

[Filter]

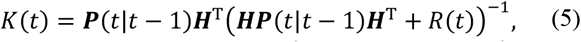

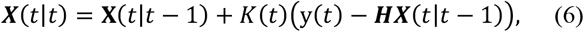

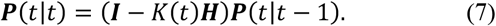

In the algorithm for the one-step-ahead prediction, ***X***(*t*|*t* − 1) is the mean vector of the prediction of ***X***(*t*) and is obtained by multiplying the matrix ***F***(*t*); ***P***(*t*|*t* − 1) is the variance-covariance matrix of the prediction of ***X***(*t*). In the filter algorithm, *K*(*t*) is the Kalman gain; and ***X***(*t*|*t*) and ***P***(*t*|*t*) are the mean vector and variance-covariance matrix of the filter of ***X***(*t*), respectively. The Kalman filter could be realized by repeating the one-step-ahead prediction and the filtering. Increasing the long-term prediction of the state could be realized by repeating the one-step-ahead prediction, such as ***X***(*t* + 1|*t* − 1) = ***F***(*t*)***X***(*t*|*t* − 1) (Fig. 1c). Note that the algorithm used the non-steady-state linear Kalman filter because the ***F***(*t*), *Q*(*t*), and *R*(*t*) depend on the time. These parameters are calculated based on the filtered signal for the past second on each time step, as does the conventional AR prediction method. The initial values of the Kalman filter used here were ***X***(0) = 0 and ***P***(0|0) = diag(0.1).

### 2.3. Test of real-time EEG phase prediction via the Kalman filter using recorded EEG signals

To evaluate our proposed method in a real-time situation, we tested the performance of the method by replaying the recorded EEG signals and recoding the replayed analog signal and the triggers, which were simultaneously generated based on the phase prediction method using the real-time digital signal processing system and an EEG recorder (Fig. 2). By using these devices, we could check the implementation of the real-time system without gathering new participants. The details of the recorded EEG signals are explained in subsection 2.4.

**Figure 2.**
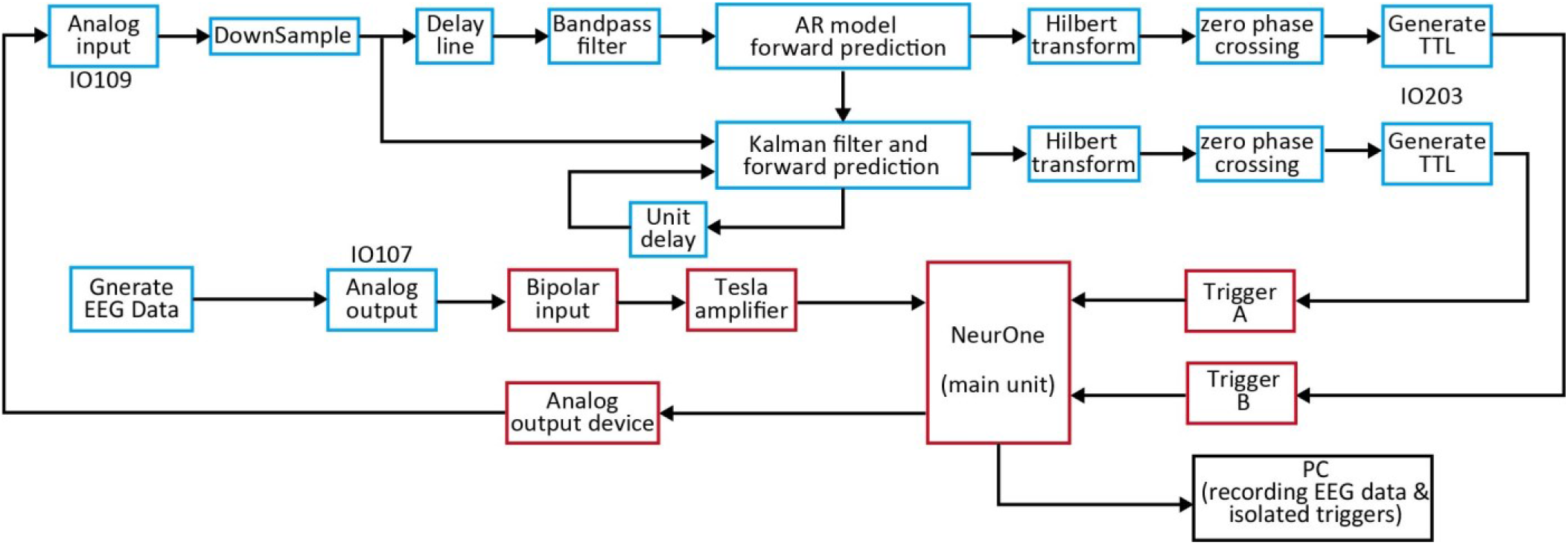
Schematic illustration of a closed-loop and online EEG replaying system. The system consisted of two main units: the performance of a real-time target machine and the NeurOne EEG amplifier system.

The system used in EEG replaying experiment consisted of a biosignal amplifier (NeurOne Tesla with Analog Out Option, Bittium Biosignals, Ltd., Kuopio, Finland) and a real-time signal processing system. The amplifier was used for recoding the artificial EEG signal, and data were acquired in AC mode with a head-stage sample rate of 20 kHz for subsequent analysis. The artificial EEG signal was re-recorded as a bipolar input. The recorded signal was sent to the real-time digital signal processing system through the analog output device. The analog output device of the amplifier was configured to recreate EEG signal from the digital data, which were filtered and amplified parallel analog signals from a user-selectable subset of 16 amplifier channels.

The real-time digital signal processing system consisted of a dedicated xPC target PC running the Simulink real-time operating system and IO devices (Performance real-time target machine SN4200, IO107, IO109, and IO202, Speedgoat, Ltd., Bern, Switzerland). The system can simultaneously generate the EEG signal and predict the EEG phase in real-time using our method via the Simulink real-time model (MathWorks, Ltd., Natick, MA, USA). The artificial EEG signal was sent to the NeurOne amplifier through the analog output device of the real-time system (IO107). The EEG data were again acquired from the NeurOne analog output device using a data acquisition card (IO 109) at a sample rate of 10 kHz, a range of ±10 V, and a 16-bit resolution. The sampling rate of received data was changed to 500 Hz. EEG phase prediction was performed by two methods: the conventional AR model prediction and our proposed method using the Kalman filter. Based on the estimated phases, two types of TTL triggers were generated by a TTL generating device (IO202) at the EEG phase of 0, which corresponds to the peak. These TTL triggers were sent to NeurOne as isolated triggers A and B, respectively. The real-time programs were asynchronously controlled from a standard PC running Microsoft Windows 10 and MathWorks Matlab R2018a through an Ethernet connection.

### 2.4. EEG data

We used real EEG data, which have been used in previous studies for different purposes [21, 27–29]. The EEG data were used in an offline analysis and the artificial EEG signal generation experiment, in which the recorded EEG signals were played and processed. The experiment from which EEG data were obtained recruited 21 participants (11 female; mean age, 26.2 years; standard deviation, 7.1 years), who provided informed consent before starting the experiment. Participants were asked to rest and close their eyes during the 3 minute EEG recording. The experiment followed the tenets of the Declaration of Helsinki and was approved by the ethics committee of RIKEN (approval no. Wako3 26-24).

For EEG recordings, an EEG amplifier (Brain Amp MR+, Brain Products GmbH, Gilching, Germany) was used with a 63-channel EEG cap (Easycap, EASYCAP GmbH, Herrsching, Germany). The sampling rate of EEG recordings was 1000 Hz, and the low and high cutoff frequencies of the online filter were 0.016 and 250 Hz, respectively. Electrodes were placed over the scalp according to the 10/10 system (AFz as the ground electrode; the reference electrode on the left earlobe). Before analyzing the data, we changed the reference to the average signal of the right and left earlobes and downsampled it to 500 Hz. All analyses were performed using custom-written scripts in Matlab (MathWorks, Inc., Natick, MA, USA) and EEGLAB. EEG signals of all 63 electrodes were used in the offline EEG phase prediction analysis. In the experiment of replaying the EEG signals, EEG data from electrodes O1 and O2 only were used.

### 2.5 EEG analysis

We calculated the four following indices to evaluate the prediction performance: time-averaged alpha power, EEG phase prediction accuracy, phase-triggered response (PTR), and phase-locking factor (PLF).

In the offline analysis, we calculated the time-averaged alpha power and accuracy of phase prediction. First, the time-varying alpha power of the EEG signal was calculated using the wavelet transform to evaluate the fluctuation of alpha oscillations. We chose the Morlet (Gabor) wavelet as the mother wavelet, which was defined using the following equation [30]:

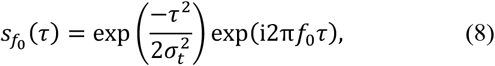

where 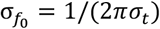, τ is time, *f*_0_ is the central frequency, and σ_*t*_ is the standard deviation of the Gaussian window in the time domain. The 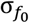 is expressed by σ_*t*_, which is the standard deviation in the frequency domain around the central frequency *f*_0_. The ratio of the frequency to variance, which characterized the mother wavelet, was set as *f*_0_/σ_*f*_ = 5. The wavelets are normalized by 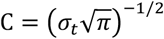, with the total energy of the wavelet being 1. By convolving the mother wavelet to the EEG time-series *y*(*t*), the time-varying EEG power was computed using the following equation:

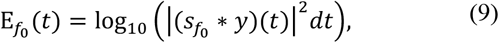

where ∗ indicates the convolution operator, E_*f*_ is the time-varying power at the central frequency *f*_0_, and *dt* is the size of the time step. To obtain an index of the stability of alpha oscillations, we calculated the time-averaged alpha power using the following equation:

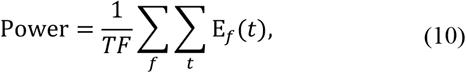

where *T* is the number of samples, *F* is the size of *f*, and the range of *f* is 8–13 Hz in 1 Hz steps.

To evaluate the accuracy of phase prediction, we defined the correct phase *ϕ_filter_* for the conventional AR prediction and Kalman filter method as the phase calculated from bandpass-filtered and Kalman-filtered signals using future EEG data, respectively (Fig. 3a). The predicted phase *ϕ_predictico_* was calculated from past EEG data only. Here, the current time was set as 0 ms, as shown in Figures 4 and 5. The phase differences defined as *ϕ*_*filter*_ − *ϕ*_*predictico*_ were calculated, and the accuracy of phase prediction was defined as the ratio between the number of phase differences, which were between π/4 and −π/4, divided by the number of all trials. By using these two indices, we confirmed the relationship between the time-averaged alpha power and the prediction accuracy of the AR prediction method.

**Figure 3.**
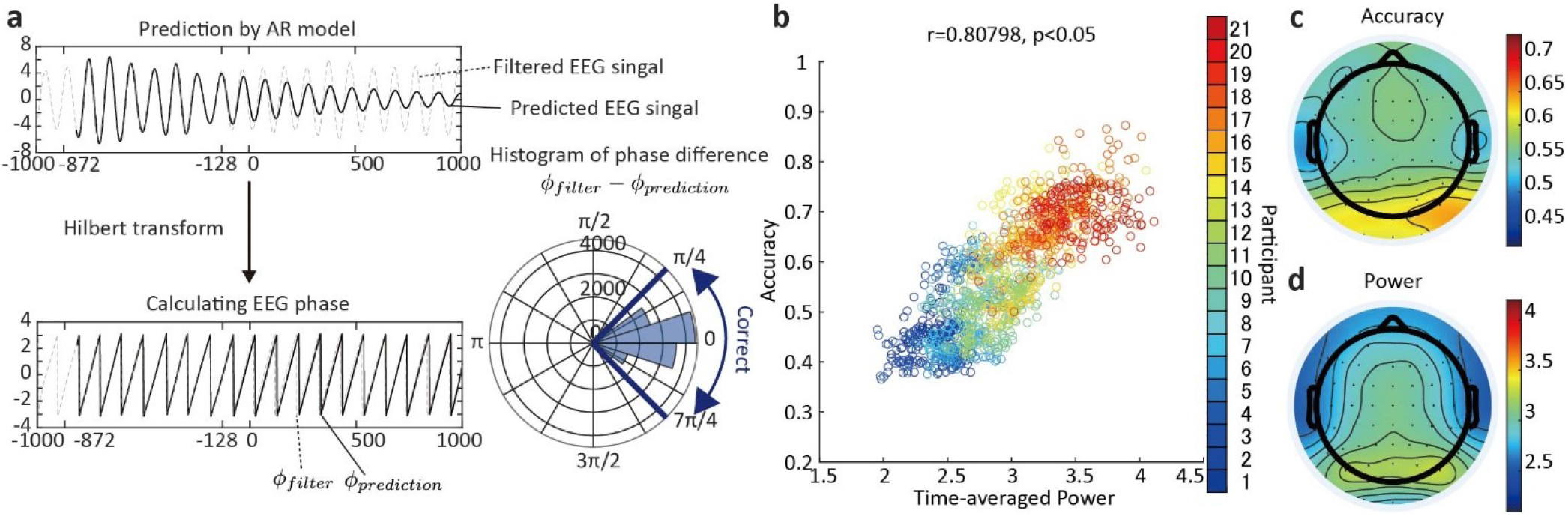
The relationship between alpha power and prediction performance for the conventional method. (a) Assessing the accuracy based on the phase difference between filtered and predicted signals. (b) The relationship between the alpha power and accuracy of prediction. The plot shows the power and accuracy of signals from all electrodes in all participants. The same color indicates the results of the same participants, including all different electrodes. (c) The topography represents the average accuracy. (d) The topography represents the average alpha power (8–13 Hz).

**Figure 4.**
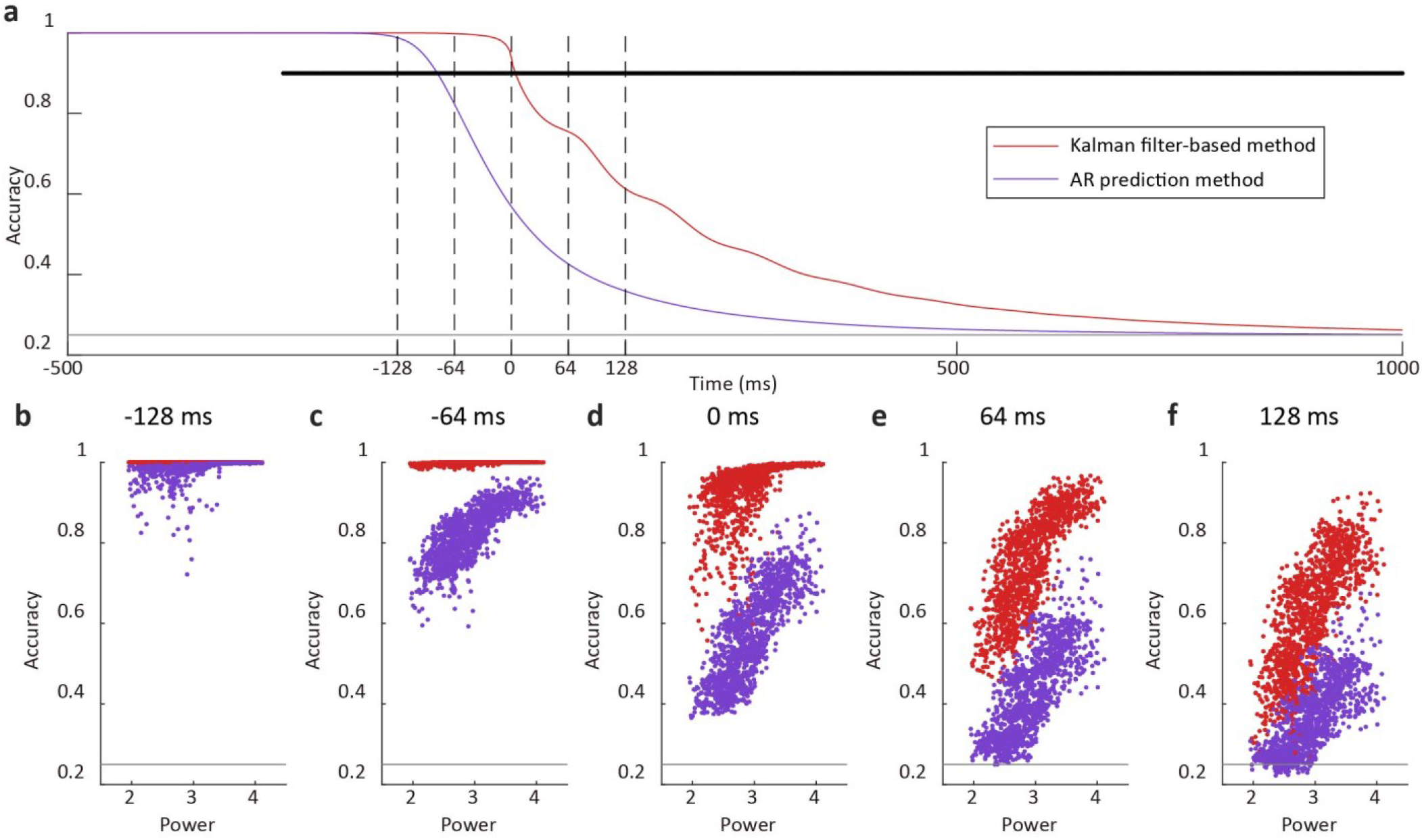
Comparing the prediction performances of the Kalman filter-based method and AR prediction method. (a) Each colored line represents the average accuracy for the Kalman filter-based method and AR prediction method. The gray horizontal line represents chance level. The black dots represent significant differences based on the Wilcoxon test (p < 0.05). (b) The scatter plot shows four types of distributions for each accuracy and alpha power at −128 ms, at which point prediction of the future signal began for the AR prediction method. (c) The distributions of accuracy and alpha power at - 64 ms. (d) The distributions at 0 ms when this was the starting point for prediction of the future signal using the Kalman filter-based method. (e) The distributions at 64 ms. (f) The distributions at 128 ms.

**Figure 5.**
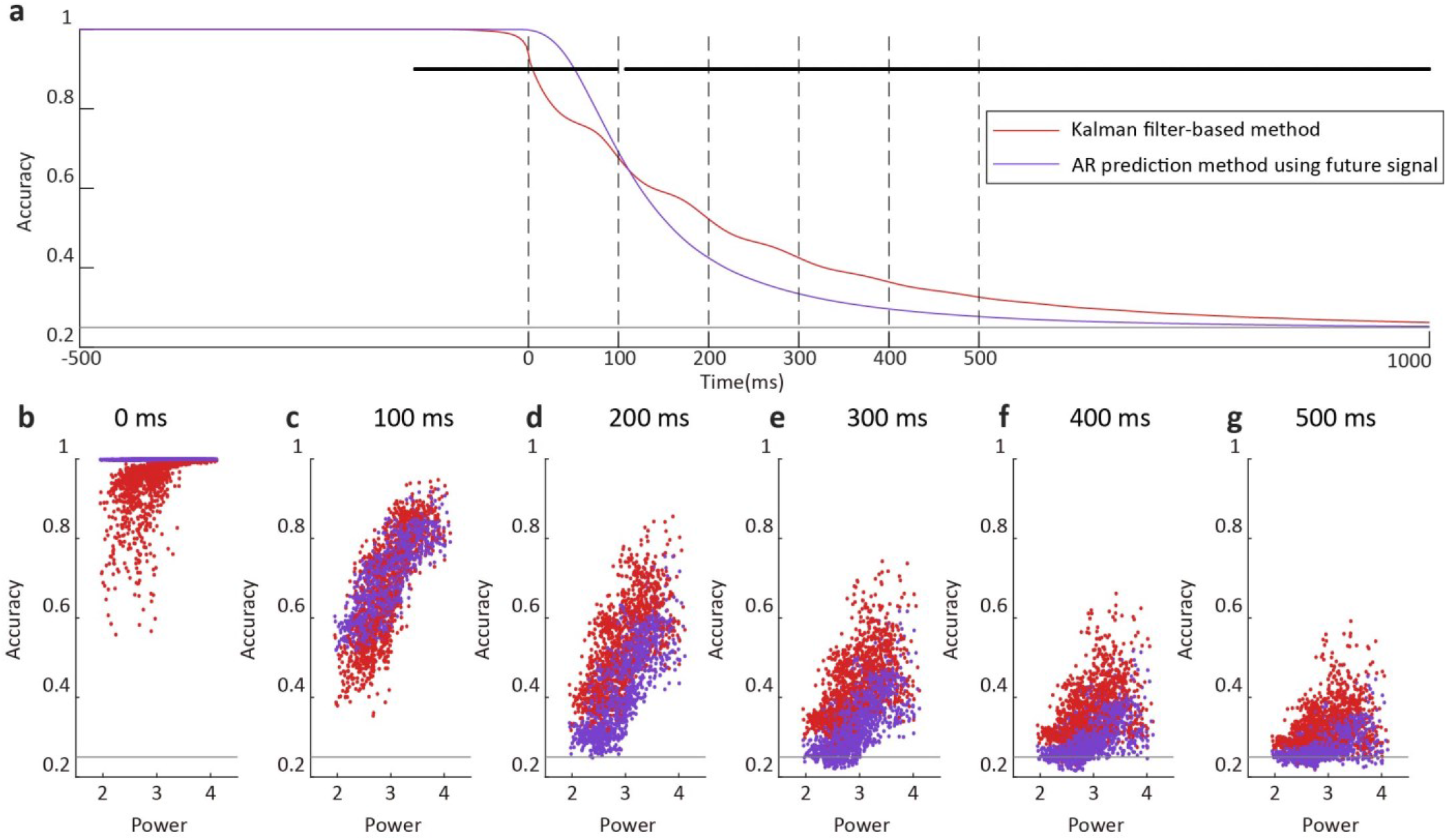
Comparing the prediction performance removing the advantage of starting point of prediction. (a) The red line represents the average accuracy based on the Kalman filter-based method, which is the same as the results shown in Figure 4. The purple line represents the average of accuracy based on the AR prediction method using future signal and used the filtered signal until 0 ms. (b) The scatter plot shows the distributions of accuracy and alpha power for the Kalman filter-based method and AR prediction method using future signal at 0 ms, which is the starting point for prediction of the future signal. (c) The distributions at 100 ms. (d) The distributions at 200 ms. (e) The distributions at 300 ms. (f) The distributions at 400 ms. (g) The distributions at 500 ms.

To confirm the results of the experiment of replaying EEG signals, we calculated the PTR [22] and PLF [30]. The PTR was defined as the grand-average of triggered EEG signals and was calculated using the following equation:

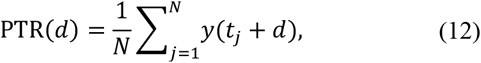

where *t*_*j*_ is the index of the sampling timepoint at which the EEG phase-dependent trigger was received, and *N* is the total number of trials. *d* is the sampling index for the PTR and ranges from −500 to 500 samples, which corresponds to the time range (from −1000 to 1000 ms relative to the stimulus onset at 0 ms). This calculation is similar to that performed for event-related potential analyses; however, it does not depend on the external stimulus and uses a generated trigger based on the EEG phase. This calculation can be used to visualize the phase-locked responses obtained by the phase prediction method when using the raw EEG signal. Therefore, this index is suitable for checking the filtering results of the EEG study.

Another index is the PLF [30], which is calculated using the following equation:

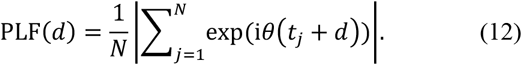

The PLF index assesses inter-trial phase locking. When the phases among trials have a high variance, the PLF is close to 0. When the phases are identical among trials, the value is 1. To obtain this index, we calculated the instantaneous phase of the alpha oscillation using the bandpass filter and Hilbert transform. The bandpass filter we used is a zero-phase digital filter with an order of 128 and a bandpass frequency of 8–13 Hz. Then, we obtained the analytic signal using Hilbert transform, as follows:

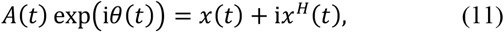

where *x*^*H*^(*t*) denotes the Hilbert transform of the bandpass-filtered signal *x*(*t*), and *θ*(*t*) is the instantaneous phase of the filtered signal [31]. *A*(*t*) is the instantaneous amplitude of the filtered signal. The same instantaneous phase calculation method for PLF was also used for the conventional AR prediction as well as our proposed method to retain a consistent estimation of the phase of alpha oscillations. The PTR and PLF were used to verify the performance of the EEG phase prediction in the real-time system.

### 2.6. Statistical comparison

To evaluate phase prediction performance differences between the conventional and proposed methods in the offline analysis, nonparametric Wilcoxon tests were applied. Specifically, we calculated p-values at each timepoint of prediction; statistical significance was defined as a two-tailed p-value < 0.05.

To compare the results of the experiment of online replaying EEG signals between the conventional and proposed methods, a cluster-based permutation test using paired t-tests [32], which allow us to avoid the multiple comparison problem, was applied to the PLF and PTR. The clusters were determined according to the temporal adjacency of the sample when it exceeded a student’s t-value corresponding to a one-tailed p-value of 0.05. The cluster-level statistics included the summation of t-values within a cluster. To determine the cluster-level threshold, the null distribution was generated by randomly shuffling the labels for the conventional and proposed methods within each individual. The procedure was repeated 10,000 times, and the maximum cluster-level statistics were computed to obtain the empirical null distribution. The statistical threshold was decided by the empirical distribution of the maximum cluster-level statistics (p < 0.05).

## 3. Results

### 3.1. The relationship between phase prediction accuracy and time-averaged alpha power for eyes-closed resting EEG data

We characterized the predictability of the EEG phase in relation to the time-averaged alpha power using the conventional method. The results revealed that the time-averaged alpha power can explain the accuracy of phase prediction very well (Pearson’s correlation coefficient *r* = 0.808, p < 0.05; Fig. 3b). The scatter plot in Figure 3 shows the phase prediction results based on EEG signals from 63 electrodes for all 21 participants. The topographies show the average accuracy at the current time (0 ms) and the average alpha power (Fig. 3c and d). These results are consistent with the previous study [16], and the relationship between the accuracy and time-averaged EEG power allowed us to visualize and evaluate the phase predictability of each recorded EEG signal. Therefore, we tested the prediction performance and time-averaged alpha power in an offline analysis.

### 3.2. EEG predictability using a Kalman filter

To test the performance of our proposed method, we applied the two prediction methods to eyes-closed resting EEG data: the conventional AR prediction method and our proposed Kalman filter with the AR model method (Fig. 4). The accuracy averages of both conventional AR prediction and Kalman filter methods sharply decreased at the prediction starting points, at which are −128 and 0 ms, respectively (Fig. 4a–b and d), and the average accuracy of the Kalman filter method continued to exceed that of the conventional AR prediction method until up to 1000 ms. These results indicate that our proposed method had a strong advantage related to the difference in the starting point of prediction.

To test the prediction performance excluding the difference in the starting point of prediction, we used the conventional AR prediction method with bandpass-filtered signals until 0 ms using the future signal (Fig. 5). The analysis completely avoided the edge effect of bandpass filtering caused by the zero paddings using the future signal. We compared the result of the AR prediction method using future signals with that of the Kalman filter method to assess the effect of using the Kalman-filtered signal on the ability to predict the future signal. The performance of the AR prediction method using future signals was significantly higher than that of the Kalman filter method from 0 to 100 ms (Fig. 5a–c). However, after 100 ms, the performance of the Kalman filter method overtook that of the AR prediction method using future signals (Fig. 5a and d–g). This suggests that using a Kalman-filtered signal is more suitable than using a bandpass-filtered signal for long-term prediction.

### 3.3 Real-time EEG phase prediction

Finally, we ran online experiments in which we replayed recorded EEG signals and real-time phase prediction (Fig. 2) to evaluate whether our proposed method (Kalman filter method) can work well for real-time EEG data. For this, we used real EEG data recorded from O1 and O2 electrodes.

The PTR amplitudes for the Kalman filter method were larger than those of the AR prediction method at 0 ms (Fig. 6a–b). The PTR allowed us to evaluate the position at which triggers occur because it only uses trigger information and raw EEG signals. Therefore, these results suggest that the Kalman filter has the same filtering performance as the bandpass filter to calculate the PTR, and a better prediction performance than the conventional AR prediction method (cluster-based permutation test, p < 0.05).

**Figure 6.**
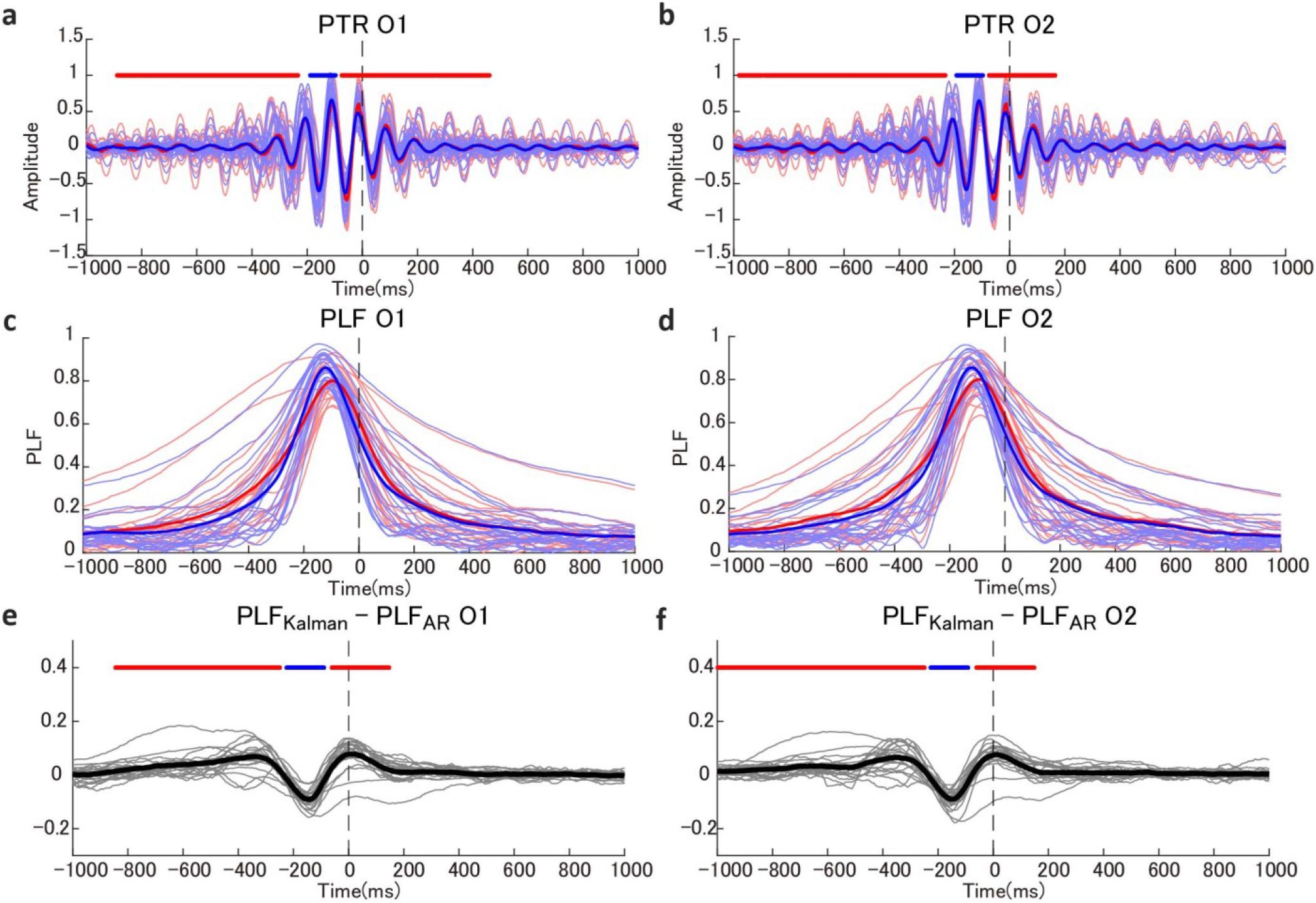
The PTR and PLF results in a real-time phase detecting system. (a) The PTR for the Kalman filter-based method (red line) and AR prediction method (blue line) using EEG signals recorded from electrode O1. The bold lines represent the average PTR. The blue and red horizontal lines represent the results of the cluster-level statistical test (p < 0.05). (b) The PTR for the Kalman filter-based method (red line) and AR prediction method (blue line) using EEG signals recorded from electrode O2. (c) The PLF for the Kalman filter-based method (red line) and AR prediction method (blue line) using EEG signals recorded from electrode O1. (d) The PLF for the Kalman filter-based method (red line) and AR prediction method using EEG signals recorded from electrode O2. (e) The differences between the PLF for the Kalman filter-based method and AR prediction method using EEG signals recorded from electrode O1. The blue and red horizontal lines represent the results of the cluster-based permutation test (p < 0.05). (f) The differences between the PLF for the Kalman filter-based method and AR prediction method using EEG signals recorded from electrode O2.

The PLF of the Kalman filter method was higher than those of the AR prediction method around and after 0 ms (cluster-based permutation test, p < 0.05; Fig. 6c–f). The results of PLF indicate that the variance of phase among all trials based on the Kalman filter method was lower than the variance based on the conventional AR prediction method. However, the PLF value of the AR prediction method was higher than the PLF of the Kalman filter method before 0 ms. The trigger of the Kalman filter method did not depend directly on the zero-phase bandpass filter. Therefore, the AR prediction method was better than the Kalman filter method before −128 ms, which was the starting point of prediction.

## 4. Discussion

Our proposed method for closed-loop EEG experiments accurately predicted the future EEG phase by introducing the Kalman filter in a real-time condition. We confirmed the improvement of our proposed method by comparing it with the conventional AR prediction method [16, 19] in an offline analysis and online experiments that involved replaying recorded EEG signals in real-time conditions. This comparison showed that our proposed method performed better than the conventional AR prediction method. Our proposed method was based on the conventional AR prediction method [19] with the addition of a Kalman filter. Notably, the prediction method of the Kalman filter, whereby the one-step-ahead prediction is repeated, is the same as that of the conventional AR prediction method. However, using the Kalman-filtered signal resulted in better performance and following significant advantages.

One advantage of our method is that it avoids the problem of the trade-off between the performance of the bandpass filter and the length of the prediction interval [16]. The conventional AR prediction method uses the bandpass-filtered signal to predict the future signal. In this case, the edge data, which are influenced by zero-padding, needed to be excluded. Therefore, to detect the current EEG phase using the conventional AR method, it is necessary to predict the long interval, which includes the excluded signal intervals. Conversely, the Kalman filter method allows us to obtain the current filtered signal without excluding the edge data. This performance of phase prediction is largely improved.

In addition, our results revealed that using the Kalman-filtered signal conveys another advantage in that it can predict more of the future EEG signal (Fig. 4). To identify advantages other than the prediction length, we compared the Kalman filter method and AR prediction method using future signals, which does not require the exclusion of the edge data due to its use of the future EEG signal. Note that the AR prediction method using future signals does not work in real-time conditions due to its use of the future signal. Although the AR prediction method using future signals has a large advantage in this way, the prediction performance of the Kalman filter method was higher than the AR prediction method using future signals after 100 ms. We suspect that the inconsistency between the filter and prediction causes a decrease in prediction performance.

Furthermore, as a result of avoiding the problem of trade-off, it is not necessary to nervously select the value of parameters in the bandpass filter in our proposed method. The trade-off between the performance of the filter and the length of the prediction interval makes the conventional AR prediction method difficult to choose the parameter [16]. The problem is caused by the edge effect of the zero-phase bandpass filtered signal. By contrast, our Kalman filter-based method does not have the trade-off, because the method used the filtered signal, which is estimated by the Kalman filter. Therefore, our method does not require sensitive parameter selection. This is an advantage when implementing closed-loop EEG experiments.

We evaluated the performance of the prediction of EEG phase using time-averaged EEG power. The time-averaged EEG power depends on the frequency of alpha oscillations during the eyes-closed resting task and the magnitude of time-varying EEG power. A previous study found that there is a relationship between the SNR, which is the ratio of targeted EEG power among the sum of powers for other frequencies; the EEG amplitude; and the phase prediction performance at the current time point [20]. Our results also revealed this relationship and additionally demonstrated the temporal changes in the prediction performance. In particular, the performance of our method using the Kalman filter decreased from the starting point of prediction and closes, before slowly reaching chance level. This temporal change might indicate that the individual EEG data comprise those of both predictable and unpredictable trials. The initial decrease might reflect the presence of unpredictable trials caused by noise or nonstationary EEG dynamics. Conversely, the performance of the conventional method decreased smoothly, which might be a result of using the non-causal bandpass filter whose order is 128, which corresponds to 256 ms. Furthermore, our results also suggest that using EEG power to determine the threshold for generating the trigger, as in a previous study [16], is an effective approach to maintain high performance and decrease individual differences in prediction performance.

In this study, we tested our proposed method in the real-time replaying of recorded EEG signals generated from an analog output device. The results showed that our method can accurately predict the EEG phase in real-time. This approach of replaying recorded EEG signals was technically useful in developing our novel method and implementing it in real-time because it allowed us to rerun the online experiment in a repeatable way. Our study not only contributes to the development of a method to reveal the role of ongoing neural oscillations, but also contributes to the development of brain– computer interfaces for clinical studies. For instance, use of the closed-loop method with deep brain stimulation has been associated with clinical improvements in patients with Parkinson’s disease [33–35]. However, in the clinical situation in which the invasive stimulation is used, it is not generaly easy to use a novel approache due to the risk such as safety. For this problem, our real-time framework without human participants by replaying recorded EEG signal is effective for testing the novel approach.We, therefore, expect that our framework to evaluate the closed-loop approach without using participants will help develop novel brain-computer interfaces for clinical use. In addition, it should be noted that the real-time framework without human participants is useful under the COVID-19 pandemic situation because we can avoid unnecessary contact with human participants.

## 5. Limitations

For our Kalman filter-based method, it is not necessary to manually select the value of parameters; however, the parameters used here were not optimal for predicting the EEG phase. If exact optimal parameters are necessary, we should perform a maximum likelihood estimation using a numerical optimization method. Numerical optimization generally requires considerable computation, and the numerical optimization method in real-time is not possible with the real-time closed-loop method considered here. If the maximum likelihood estimation is necessary for offline analysis, we suggest using the parameters decided here as initial values of the numerical optimization.

In this study, we did not consider colored noise, which includes 1/f fluctuation, or the effect of TMS, eye-blink, and muscle artifacts. If noise and artifacts can be defined, the Kalman filter will be able to remove them from the raw EEG signal by including the model of them. In fact, there are some methods using the Kalman filter assuming colored noise [36] and excluding artifacts by the Kalman filter in an offline analysis [37, 38]. We did not use a complex model to avoid increasing the number of parameters and instead aimed to reveal the effect of simple Kalman filtering. However, the removal of noise and artifacts using a Kalman filter might be an effective approach to develop an automated artifact rejection method in real-time to improve phase prediction accuracy.

To apply our method to closed-loop EEG experiments, some problems remain. For instance, parameters of individual participants, such as the target time of prediction and the electrode signal used to make the prediction, should be investigated case by case. Future work should focus on how to detect the parameters relevant to each participant before or during the experiment.

## 6. Conclusions

We proposed a novel phase-dependent stimulation method using the Kalman filter and constructed a framework to evaluate the method in real-time. This study contributes to the development of an effective method for closed-loop EEG experiments and their implementation.

## Acknowledgments

The project was supported by a research grant from Toyota Motor Corporation, the Special Postdoctoral Research Program at RIKEN, and JSPS KAKENHI Grant Numbers 19K21120 and 20K19887.

